# Localizing the Sources of Modulations of the P3m Component by Task Difficulty

**DOI:** 10.1101/2022.09.09.507266

**Authors:** Cindy Boetzel, Florian H. Kasten, Christoph S. Herrmann

**Affiliations:** Experimental Psychology Lab, Department of Psychology, European Medical School, Cluster for Excellence “Hearing for All”, Carl von Ossietzky University, Oldenburg, Germany; Neuroimaging Unit, European Medical School, Carl von Ossietzky University, Oldenburg, Germany; Research Center Neurosensory Science, Carl von Ossietzky University, Oldenburg, Germany; Centre de Recherche Cerveau & Cognition, CNRS, Toulouse, France; Université Toulouse III Paul Sabatier, Toulouse, France

**Keywords:** P300, P3m, Sources, Event-related fields (ERF), task difficulty, Magnetoencephalogram (MEG), P3m Modulation

## Abstract

The P3m is a late component of the event-related field (ERF) and is assumed to reflect the allocation of resources in various cognitive domains. The amplitude of the P3m can be affected by several task parameters, like stimulus probability, inter-stimulus interval and task difficulty. The task difficulty can alter the P3m amplitude, depending on which mechanisms are involved. The sources of the P3m are located in several brain areas, suggesting a widespread network generating the P3m. Which of these sources of the P3m give rise to the task-difficulty induced amplitude modulation are not well investigated. However, localizing these sources could pave the way for intervention studies with non-invasive brain stimulation methods, which might be suitable to modulate the pathologically altered P3m emerging in various psychiatric and neurodevelopmental disorders. We designed a MEG-study with a visual oddball-like task with two different task difficulties to 1) show decreased P3m amplitudes induced by increased task difficulty and 2) estimate the sources of this P3m modulation. Additionally, the influence of the increased task difficulty on behavioral outcomes were analyzed. The P3m amplitude decreased significantly in the hard condition compared to easy condition. Furthermore, the hit rate for standard trials decreased significantly, while the reaction times increased for both, targets and standards. For this specific visual oddball-like task, the sources of the task difficulty dependent P3m amplitude modulation are located in the centro-parietal regions.

## 2 Introduction

The P3m, whose counterpart in the electroencephalogram (EEG) is the P3, is a late component of the event-related field (ERF) evoked by events of various modalities. Typically, the P3m occurs 300 – 600 ms after stimulus onset and is characterized by a large positive deflection (Luck, 2014), which can be influenced by different task parameters like stimulus probability (Polich, 1990), inter-stimulus interval (Polich, 1987) and task difficulty (Hagen et al., 2006; Kim et al., 2008; Selimbeyoglu et al., 2012). The task difficulty is negatively correlated with the amplitude of the P3m. But it is hitherto unknown which brain areas are involved in the task difficulty dependent modulation of the P3m amplitude?

Several studies investigated where the visual P3m originates. For the P3m, various inconsistent source locations were reported. For example, Rogers et al., (1993) identified two simultaneous but spatially distinct P3 sources. One in deep structures of the right hemisphere under the temporal cortex, near the right hippocampal formation and the second one in the primary visual cortex (Rogers et al., 1993). Okada et al., (1983) revealed the source of the visual N2-P3 complex in the hippocampal formation and Mecklinger et al., (1998) identified neuronal generators placed in subcortical structures in the vicinity of the thalamus. Next to these sources, the superior and inferior frontal lobe, the middle temporal gyrus, the parietal lobe and the cingulate gyrus were shown to represent prominent sources of the P3m (Sabeti et al., 2016). These results give evidence for a widespread network involved in the P3m generation. To the best of our knowledge, no study showed the locations of the P3m amplitude modulation affected by task difficulty. However, a clear identification of the sources of the P3m and the associated modulation of the P3m by task difficulty in the specific modality and task could pave the way for intervention methods like non-invasive brain stimulation methods (NIBS). NIBS might be suitable to modulate the pathologically altered P3m emerging in several psychiatric and neurological disorders (Hilger et al., 2020; Määttä et al., 2005; Neuhaus et al., 2010; Wiersema et al., 2006). Therefore, the location of the P3m amplitude modulation has to be identified, as for utilizing NIBS reliably as treatment method, the target regions must be clearly defined. Vaguely defined target regions could result in stimulation having little to no effect because the source(s) are not stimulated sufficiently.

Hence, we developed a visual task with two different task difficulties, which reliably evokes ERFs to investigate the sources of the task difficulty dependent P3m modulation. In a first step, we aimed to demonstrate that our task modulates the P3m amplitude. We hypothesize a decreased P3m amplitude for ERFs evoked in the hard condition compared to the ERFs evoked in the easy condition. Furthermore, we expect a reaction time (RT) increase for the hard condition (Kim et al., 2008). In the hard condition, the physical difference between target and standard stimuli is very small, hence participants need more time for the categorization and the latency and RT increase (Luck, 2014). Additionally, for the reasons stated above, we hypothesize a hit rate decrease in the hard condition as compared to the easy condition. Furthermore, we localized the task difficulty dependent P3m amplitude modulation using a Linearly Constrained Minimum Variance (LCMV) beamformer.

## 3 Methods & Material

### 3.1 Participants

For the study, 19 participants (12 females, mean age 25.4, ranging from 18 to 31) without any self-reported history of psychiatric or neurological diseases were introduced. Exclusion criteria like metallic materials within the head, the occurrence of epilepsy in the family history (1st degree) and skin diseases on the head were inquired before the experiments started. Further requirements for participation were normal or corrected to normal vision, right handedness and non-smoking. One subject was excluded due to an incorrect segmentation of the MRI-Scan, which precluded source analysis, two participants had to be excluded, because their performance in the easy condition was below 65%, one subject fell asleep while performing the tasks. Thus, 15 participants (9 females) remained for data analysis. All volunteers had to participate in a single MEG session consisting of a short training session in the beginning and two experimental blocks. Task difficulty was manipulated between blocks with the order of the conditions randomized and counterbalanced across participants. All subjects received monetary compensation for their participation. The study was approved by the “*Commission for Research Impact assessment and Ethics*” at the University of Oldenburg.

### 3.2 Visual Task

In order to evoke the ERFs, participants had to perform a visual, oddball-like task. The visual task is depicted in Figure 1 A & B. The Gabor patches, generated in Matlab R2019a (The MathWorks Inc., Natick, MA, United States), were tilted either to the left or to the right and had 30 pixels per cycle (resulting in a spatial frequency of 3.8 cycles per visual degree angle). The Gabor stimuli were presented using a custom-made Presentation script (Presentation version 18.1). The distance between the participants and the monitor was fixed to keep the perception of the stimuli constant for each participant. At the beginning of each trial, a small circle occurred at the center of the screen as a fixation point. After a randomized interval (1800 and 2800 ms), the Gabor patch occurred for 500ms at the center of the monitor. The participants had a response window of 1300ms after stimulus onset and were instructed to press the button as fast as possible. The fastest reaction time considered as correct response was set to 200 ms after stimulus onset. The participants had to respond to each stimulus by pressing the respective button (left or right) with the index finger. Each condition consisted of 400 trials and lasted ~20 minutes. The participants were assigned to one of two target directions groups. This means, when a subject was in group one, the target stimulus was tilted to the left and the standard stimulus to the right. In group two, the assignment was reversed. Target stimuli were presented with a likelihood of 25 % and standard stimuli with 75 %. This rather high target probability was chosen to ensure a sufficient number of trials for a robust ERF, while still representing an infrequent oddball-like event that should produce an increased P3M compared to standards (Kruggel et al., 2001). In order to ensure that the target stimuli were perceived as oddballs, some stimuli presentation constraints were implemented. Each condition started with a sequence of at least 10 standard stimuli and only two targets could occur consecutively, followed by a standard stimulus. The task difficulty was regulated by the tilt angle. Each participant performed a short training with three different conditions with increasing difficulties (easy: 2°; medium: 1° and hard: 0.5° from vertical line) to get familiar with the task. Each training condition comprised 15 trials, including 4 target trials. Afterwards, each participant performed two conditions (easy and hard condition) of the visual task. Participants were assigned to one of two groups, which defined the target direction for both conditions. Thus, when a participant was assigned to group one, the target stimulus was tilted to the right and the standard stimulus to the left. In group two, the assignment was inversed. The tilt direction of the target stimuli was counterbalanced across subjects.

**Figure 1.:**
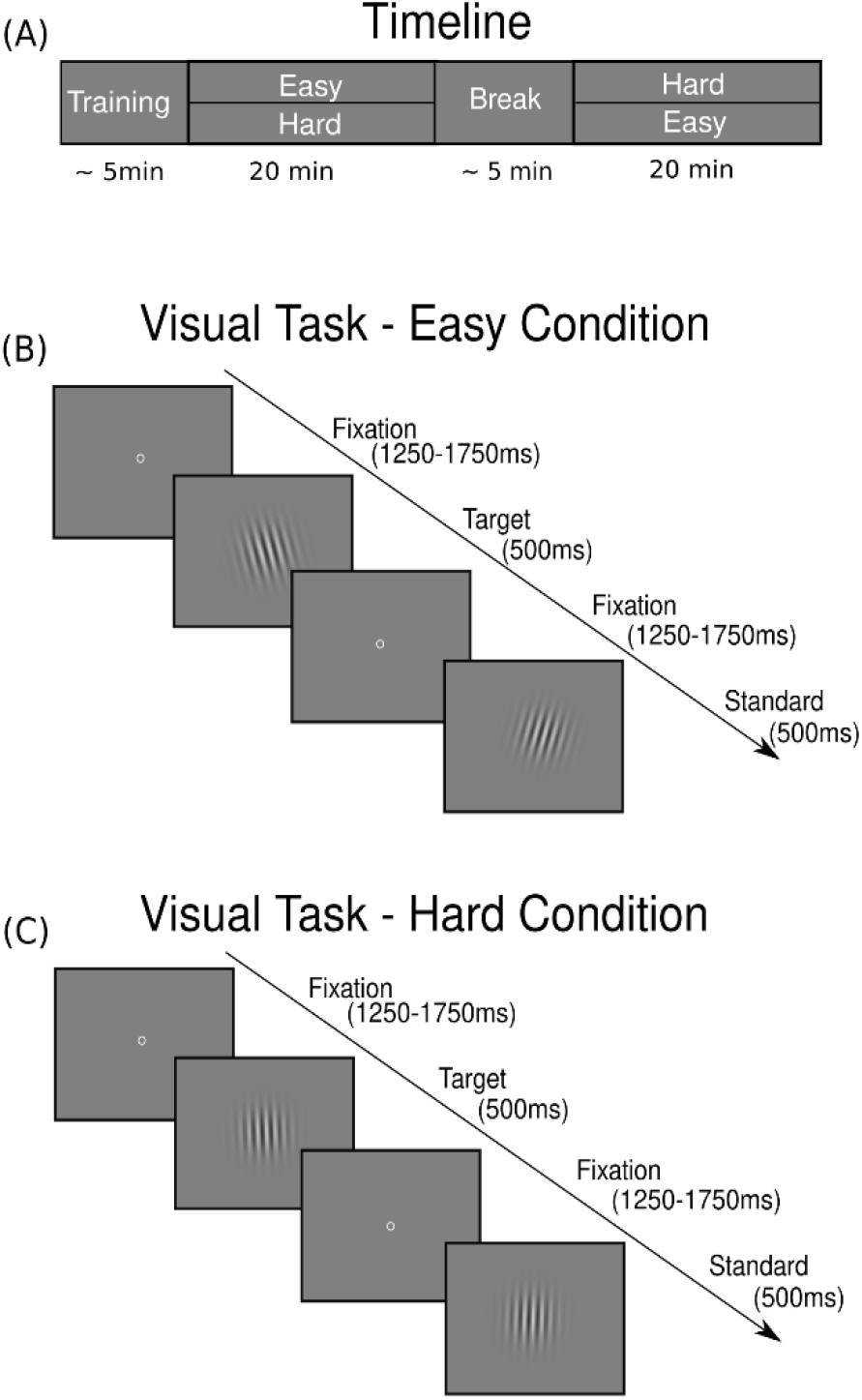
Experimental Setup. A) Timeline of the experiment. The experiment started with a short training, followed by either the easy or the hard condition. After a short break, the participants had to perform the second experimental block with the other condition. Each experimental block lasted 20 min. B) Visual task in the easy condition. Centered circles served as fixation. Inter-stimulus interval jittered between 1250 and 1750 ms. Visual stimuli occurred for 500 ms followed by an inter-stimulus interval. The tilt angle of the Gabor patch was 2° C) Visual task in the hard condition. The tilt angle of the Gabor patch was 0.5°. For visualization, the tilt angles were increased compared to the experimental tilt angles.

### 3.3 Magnetoencephalogram

During both experimental sessions, neuromagnetic signals were recorded using a whole-head 306-channel MEG with 102 magnetometers and 204 orthogonal planar gradiometer sensors (Elekta Neuromag Triux System, Elekta Oy, Helsinki, Finland). In preparation for each MEG measurement, 5 head-position indicator (HPI) coils were attached to the head of the participants - three coils at the frontal hairline and one coil at each mastoid, respectively. The position of the coils was registered together with anatomical landmarks (left and right tragus and nasion) and at least 200 samples of the head shape using a Polhemus Fastrak (Polhemus, Colchester, VT, USA). During the MEG measurements, the sensor array was in upright position (60° dewar orientation).

### 3.4 MRI Acquisition

To perform source analysis of the individual P3m modulations, a structural MRI was obtained from each subject. The MRI scans were acquired using a Siemens Magnetom Prisma 3 T whole-body MRI machine (Siemens, Erlangen, Germany). A T1-weighted 3-D sequence (MPRAGE, TR = 2000 ms, TE = 2.07 ms) with a slice thickness of 0.75 mm was used.

### 3.5 Data Analysis

The data analysis was performed in Matlab R2019a using the fieldtrip toolbox for MEG processing (Oostenveld et al., 2011). External interferences occurring in the MEG were suppressed using the spatiotemporal signal space separation method (tSSS), with standard settings (*L*_in_ = 8, *L*_out_ = 3, correlation limit = 0.98; Taulu & Simola, 2006; Taulu et al., 2005), using MaxFilter v2.2 (Elekta Neuromag, Elekta Oy, Finland). The raw data were resampled to 250 Hz. An independent component analysis (ICA) was conducted to separate the signal into additive subcomponents. Eyeblinks, eye movements and heartbeat components were removed before back-projecting the data into sensor space. Subsequently, the cleaned data were filtered between 0.1 and 10 Hz with a 4^th^-order Butterworth filter. The high pass filter was set to 10 Hz to emphasize the P3m amplitude, whose frequency ranges between the delta (0.5 – 3.5 Hz) and theta band (~ 4 Hz – 7.5 Hz; Steriade, 2003). In order to extract the ERF, the data were cut into 2s epochs (−0.5 – 1.5 s around stimulus onset) and averaged.

#### 3.5.1 Statistical Analysis

Statistical analyses were performed in Matlab R2019a using the fieldtrip toolbox. For statistical analyses, only the Gradiometer signals combined over both directions were considered. In order to investigate how the difficulty of the task influenced the P3m amplitude, the ERFs recorded by the combined gradiometers were averaged for each participant. Afterwards, a one-tailed dependent cluster-based permutation test (CBPT) with 10000 random draws (Maris & Oostenveld, 2007) and Monte Carlo estimates to calculate the p-values for data between 0 - 0.9 sec after stimulus onset was performed for each stimulus type (targets and standards) and the average of both stimulus types (combined). Time intervals exhibiting significant cluster were defined as times of interest (toi) and channel exhibiting the significant cluster were defined as region of interest (roi).

For statistical analysis of the behavioral data, the percentage of correct trails per participant was calculated. Since the Kolmogorov-Smirnov test revealed that the behavioral data were not normally distributed, a Wilcoxon–Mann–Whitney-*U* test was performed for statistical analysis.

#### 3.5.2 Linearly Constrained Minimum Variance Beamformer

The P3m amplitude modulation induced by task difficulty was projected into source-space using a LCMV beamformer (van Veen et al., 1997). The ERFs of the single subjects were calculated (from −1 to 2 s around stimulus onset). The ERF data were projected to a 6 mm grid and warped into MNI (Montreal Neurologic Institute) space. A single-shell head model for each participant was co-registered to the head position of the participant in the MEG. The ERF data were utilized to calculate a common filter which in the following was applied for the projection of the easy and hard condition data. The P3m sources were computed in a time-interval between 450 – 550 ms after stimulus onset (peak of P3m on average at 495 ms). These source data were averaged across participants for each stimulus type. In order to investigate the location of the P3m modulation by the different stimulus types and by the task difficulty, initially the difference between target and standard stimuli (in the sequel referred to as Δ stimulus) was calculated for each participant and averaged afterwards across the whole sample. For the task difficulty dependent P3m modulation representation, we calculated the difference of Δ stimulus between the hard and the easy condition (in the sequel referred to as Δ condition).

## 4 Results

### 4.1 Behavioral Data

To verify that our manipulation of task difficulty worked, we tested for changes in performance and RTs. Figure 2 A shows the hit rates for targets and standards for each block and Figure 2 B represents the RTs, respectively. Indeed, the hit rate of targets and standards decreased in the hard condition compared to the easy condition. However, only the Wilcoxon–Mann–Whitney-*U* test for the decrease of the hit rate for standard trials reached significance (Z_(14)_ = 3.941, p = < 0.01), although also a significant decrease of the hit rate was also expected for the target trials. A significant performance decrease for standard trials but not for target trials might influence the analysis of the task difficulty dependent P3m modulation. We could assume that the P3m of target trials did not decrease as strongly as the P3m amplitude of the standard trials and thus the sources of the standards weighted stronger in the difference calculation. For standards, the RTs increased significantly in the hard condition (Z_(14)_ = −2.904, p = < 0.01) compared to the easy condition, while for targets, the increase of the RT show a trend (Z_(14)_ = − 1.660, p = 0.09).

**Figure 2.:**
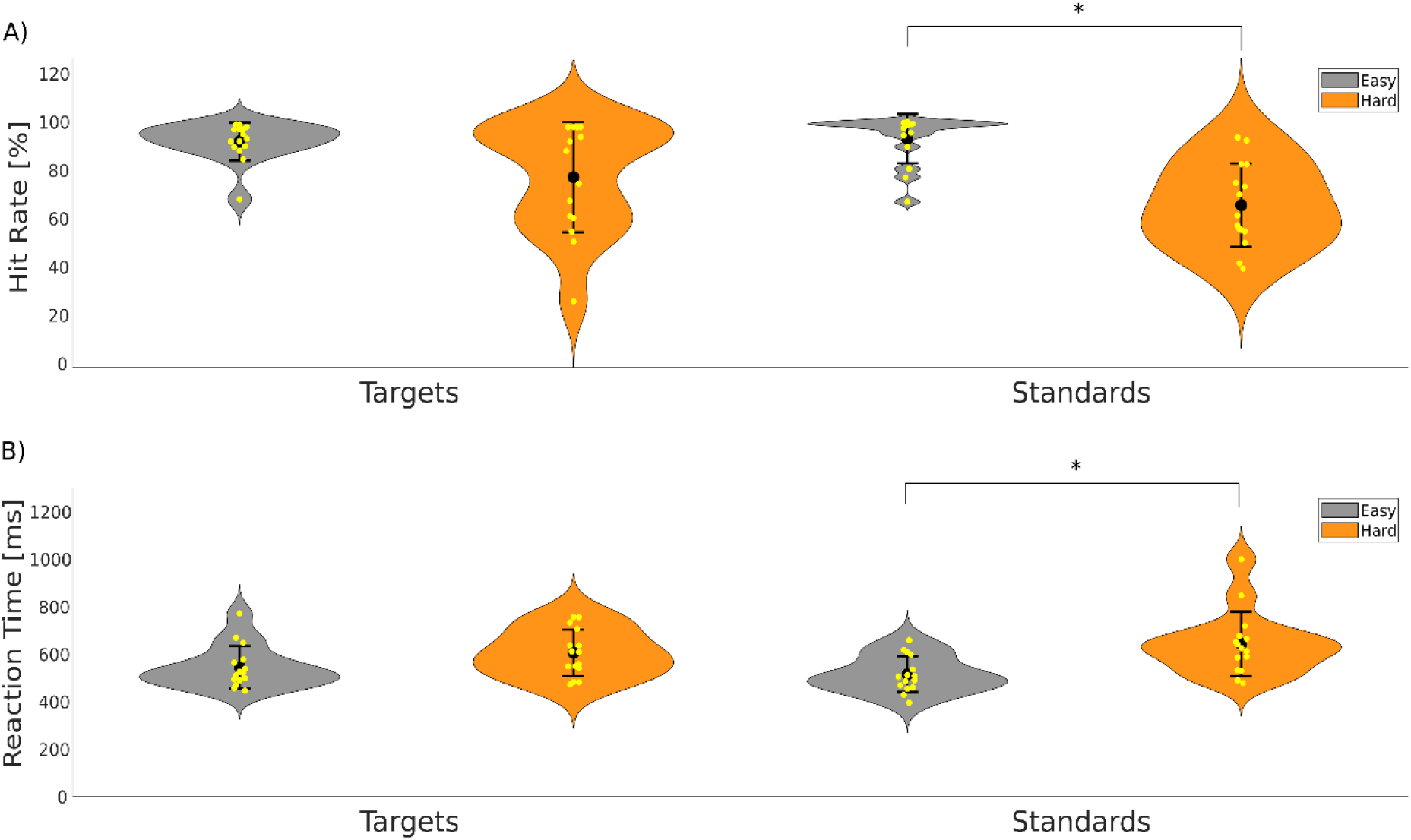
Behavioral Data. A) Hit rate for targets and standards, respectively. Yellow dots indicate single data points. Big black dot represents the mean of the data and error bars indicate the standard deviation. Grey violins represent the easy condition and orange violins the hard condition. While the violins of the targets show high overlap, the violins for the standard trials show a clear decrease in the hit rate, which is supports the statistical analysis of the data. B) RTs for targets and standards, respectively. RTs increase for both stimulus types in the hard condition, however, for targets, the violins show a slight overlap.

### 4.2 P3m Amplitude Modulation induced by Task Difficulty

We next tested if the differences in task difficulty observed on the behavioral level are reflected in the P3m amplitude on sensor level. The CBPT with an a-priori selected time window (0 - 0.9 sec after stimulus onset), revealed a difference between the easy and hard condition in each stimulus type. For targets and standards combined, there is a significant cluster ranging between ~292 – ~592 ms after stimulus onset (p < 0.01). For targets only, a significant cluster ranged between ~280 – ~600 ms (p < 0.01) and for standards between ~304 – ~584 ms (p < 0.01) after stimulus onset. Time intervals exhibiting significant cluster were defined as times of interest (toi) for further visualization steps. In order to visualize the differences between the easy and hard condition for each stimulus type in the time-domain, channels forming the significant clusters (see Figure 3, small black crosses in topographies) were averaged and the resulting signal traces are shown in Figure 3 as ERFs. The ERFs clearly show a reduced P3 amplitude in in the hard condition (red line) as compared to the easy condition (black line) for each stimulus type. The topographies show a rather fronto-central distribution of differences between the easy and hard condition, indicating that these fronto-central regions are involved in P3m amplitude modulation.

**Figure 3.:**
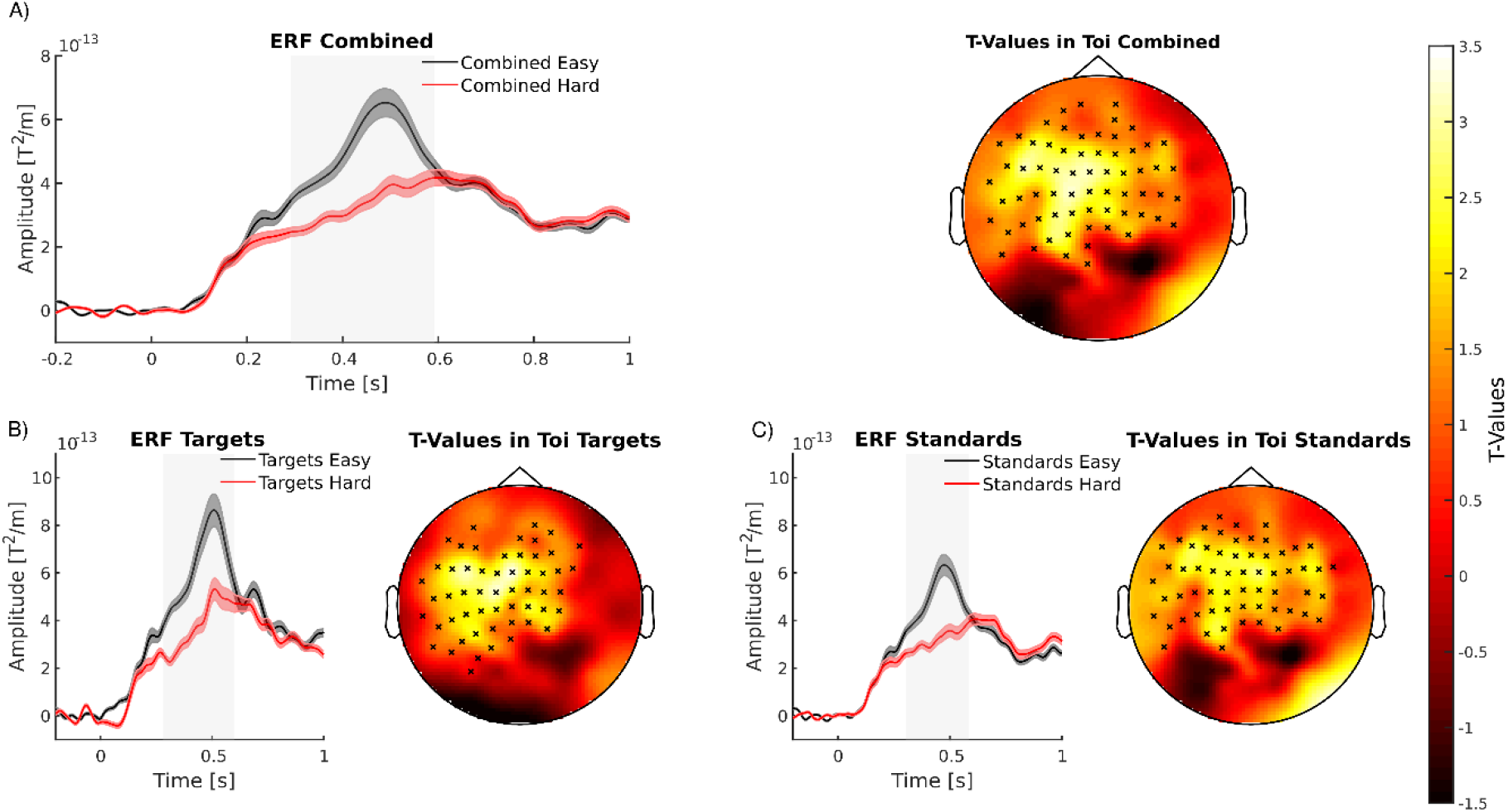
Visualization of Cluster-Based Permutation Test Results. A) Combined ERFs of gradiometers revealing a significant difference between the easy and hard condition (indicated by small crosses in topography top right). The black line indicates the target trials, red line indicates the standard trials. Shaded error bar represents the standard error. The topography (top right) shows the t-values resulted from the CBPT averaged over the toi. Black crosses indicate the channel, associated with the significant cluster. B) ERFs and topography with t-values for target trials. C) ERFs and topography with t-values for standard trials

### 4.3 Sources of Task Difficulty Dependent P3m Modulation

Averaged results of the LCMV beamformer for investigating the sources of the P3m for each stimulus are depicted in Figure 4. Figure 4 shows the absolute source activity of the gradiometers. The highest source activity for standard stimuli in the easy as well as in the hard condition is located in the centro-parietal region of both hemispheres. For target stimuli, as compared to standard stimuli, the source activity in the centro-parietal region shows higher amplitude values. Figure 5 shows the differences of the P3m amplitude modulation induced by different task difficulties. The Δ stimulus in the easy condition (Figure 5 A) is higher as compared to the Δ stimulus in the hard condition (Figure 5 B). The source activity of Δ stimulus in the easy condition as well as in the hard condition is located in the centro-parietal regions of both hemispheres. The source activity of Δ stimulus in the hard condition shows a lower amplitude and but a similar distribution, including mainly centro-parietal regions. The highest Δ condition can also be found in the centro-parietal regions.

**Figure 4.:**
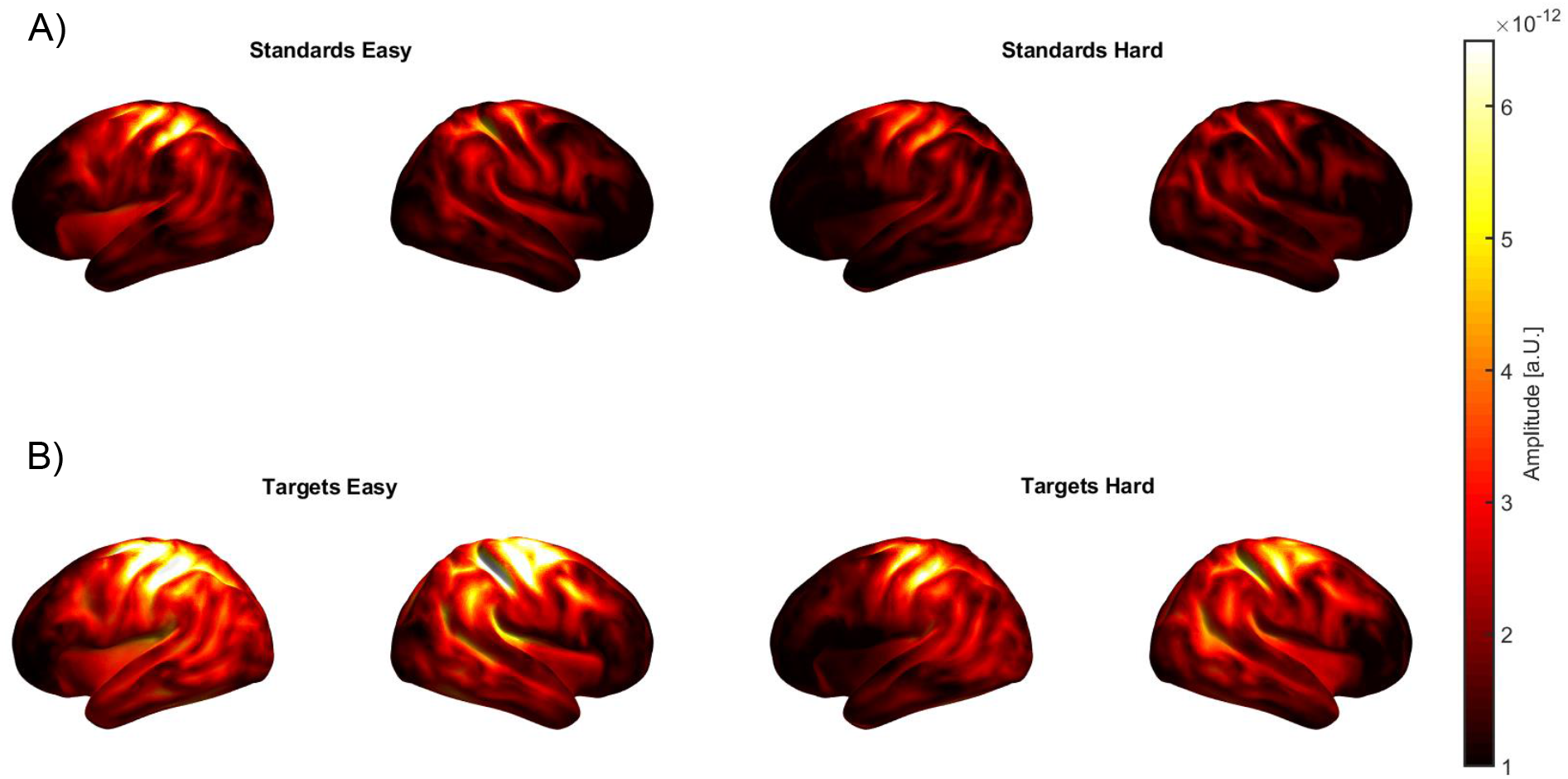
Averaged results of the LCMV beamformer for each stimulus type and each condition. A) Source activity for standard stimuli for the easy (left) and hard (right) condition. Absolute values of source activity are depicted. The highest activity for standard stimuli in the easy as well as in the hard condition is in the centro-parietal region. B) Source activity for target stimuli for the easy (left) and hard (right) condition. For target stimuli, the source activity in the centro-parietal regions is higher as compared to the source activity of standard stimuli.

**Figure 5.:**
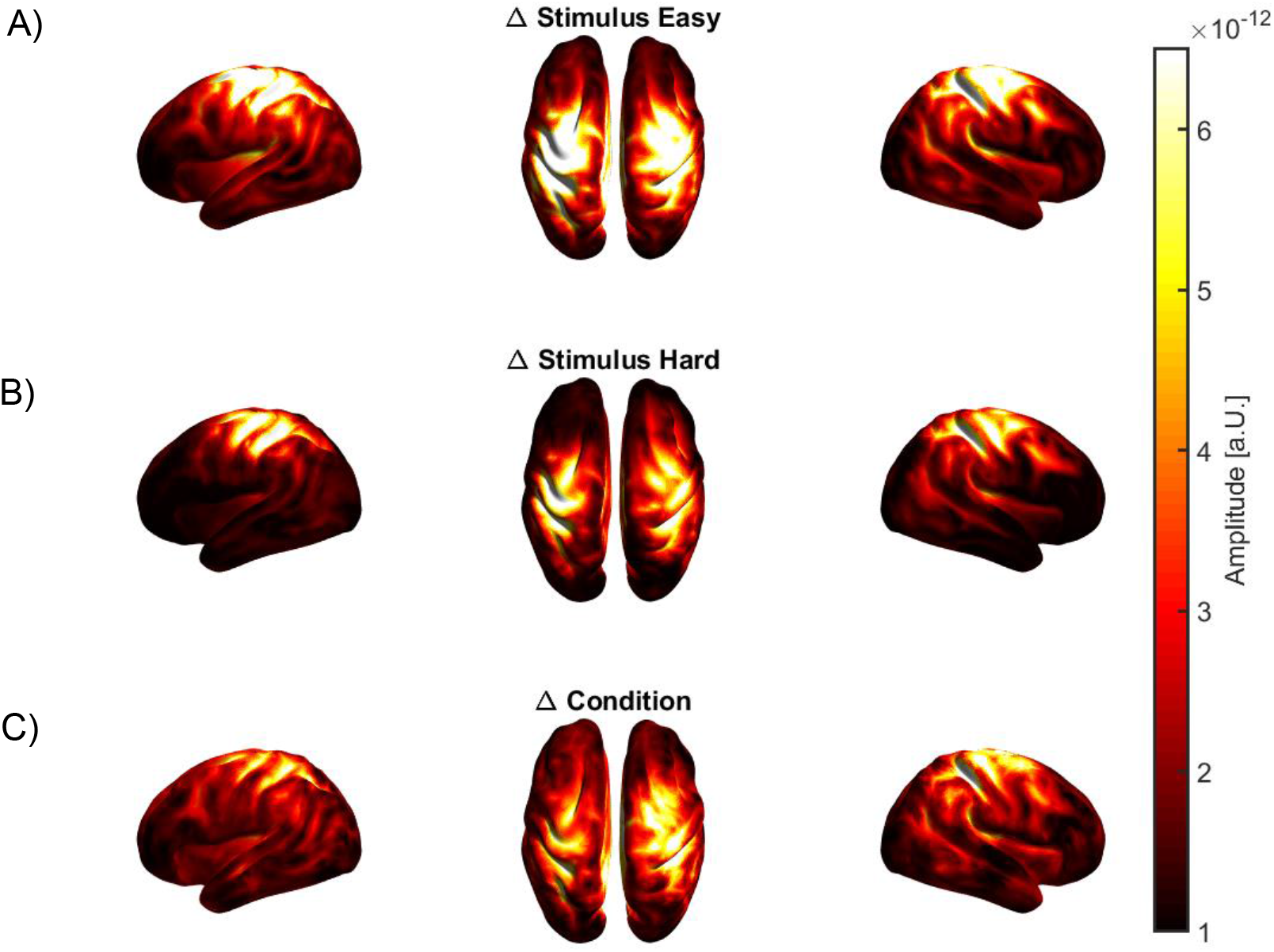
Grand averaged results of the LCMV beamformer Δ stimulus and Δ condition. A) Source activity for Δ stimulus for the easy condition, with the highest source activity in centro-parietal regions. Absolute values of source activity are depicted. B) Source activity for Δ stimulus for the hard condition with a similar distribution of source activity in centro-parietal regions but with lower amplitude values. C) Source activity for Δ condition, with highest source activity in centro-parietal regions with descriptively higher amplitude values in the right hemisphere.

## 5 Discussion

The main aim of this study was to locate the sources of the task difficulty dependent P3m modulation. Therefore, a visual oddball-like task was established, to reliably decrease the P3m amplitude as well as the performance. The modulation of the performance and the associated P3m amplitude decrease was intended to be achieved by increasing the difficulty of the visual task. We showed a decreased performance-level and increased RTs in the hard condition as compared to the easy condition. Additionally, a decrease in P3m amplitude in the hard condition could be shown. The source activity of the task difficulty dependent P3m modulation was primary located in centro-parietal regions (see Figure 5).

Previous studies demonstrated the effect of different task difficulties on the amplitude of the P3 (Hagen et al., 2006; Kim et al., 2008; Selimbeyoglu et al., 2012) with higher task difficulties lowering the P3 amplitude. These findings support the results of the present study, since the hard condition evoked a significantly lower P3m amplitude as compared to the easy condition. An explanation for these findings might be that the P3m is associated with memory demands and attentional processes (Kok, 2001; Polich, 1986; Portin et al., 2000). For the easy condition, the physical difference between the two stimulus types was designed to be such obvious that only a small amount of attention was required to distinguish the target from the standard. Additionally, the participants seemed to be certain whether the stimulus was left or right tilted. This is reflected by a high performance-level and low RTs in the easy task (see Figure 2). But with increasing task difficulty in the hard condition, the physical difference between left and right tilted stimuli was rather small. Probably the adaption of the task difficulty resulted in our study in a lower external or internal discriminability of the stimuli that the participants were less certain about the category of the stimulus. This assumption is supported by the rather high decrease in the performance-level and increased RTs (see Figure 2). This uncertainty in discriminating the stimuli is accompanied by a decreased P3m amplitude (Kok, 2001; Luck, 2014; Selimbeyoglu et al., 2012). In case the participants would have been certain about the stimulus and the classification into the correct category, but had to have allocated more attentional resources to perform the task, the P3m amplitude would have been increased (Kok, 2001; Luck, 2014). Therefore, it is reasonable to assume that the participants had greater problems classifying the stimuli into the correct category in the hard condition rather than requiring more resources to perform the task.

Further parameters which may influence the amplitude of the P3 are the inter-target interval between two stimuli, the probability of the stimuli and the relevance of the stimulus for the task (Luck, 2014). In the present study, all these parameters were kept similar between blocks to rule out these additional parameters as possible reason for differences in the P3m amplitude. Thus, we are confident that the P3m modulation is caused by the changes in task difficulty. The ERF-findings are also supported by the behavioral data, which showed a significantly decreased hit rate for standards in the hard condition as compared to the easy condition, confirming our experimental manipulation. The hit rate of the targets decreased descriptively but did not reach significance (see Figure 2). Next to a decreased performance-level, our data show increased RTs for both stimulus types in the hard condition. However, only the RT increase of standards was significant.

There is a discrepancy between our expectations and the measured behavioral data, since we also assumed a hit rate decrease for target stimuli, which could not be observed in our data. Here, it has to be considered that this paradigm is an oddball-like paradigm, thus the target stimuli are much less frequent (25%) than the standard trials (75%). Therefore, by the number of trials alone, the decreasing effect of the performance-level is probably much better estimated for the standards compared to the targets. Note that, descriptively, the hit rate for stimulus types decreases in the hard condition.

As already mentioned above, the sources of the P3m are widely distributed and a widespread network seems to be involved in P3 generation (Sabeti et al., 2016). For possible future applications of NIBS aiming to modulate the ERF and especially the P3m, the distributed sources pose an obstacle, because stimulating only vaguely defined sources will have little to no effect. Hence, defining clear sources is a prerequisite for stimulation effects. The present results of the LCMV beamformer show source activity in both centro-parietal regions for target and standard stimuli and for Δ condition. These findings go in line with the results of several studies investigating the P3m sources in the visual domain (for a review considering the visual and auditory modality, see Linden, 2005). For example, Bledowski et al., (2004) showed in a functional magnetic resonance imaging (fMRI) study P3b sources in the inferior parietal lobe, posterior parietal cortex and inferior temporal cortex for participants performing a difficult visual three-stimulus oddball paradigm. Furthermore, Halgren et al., (1995) identified superior parietal regions in intracranial recordings in epileptic patients to contribute to the P3b generation. Another study, conducted by Prabhu et al., (2001) in which the participants (alcoholics and healthy controls) had to perform a visual-oddball task the P3 source activity of the visual P3 was localized in bilateral prefrontal and central regions (Prabhu et al., 2001). Next to P3 sources in the parietal regions, further brain areas were identified to be active, when a P3 is evoked. For example, McCarthy et al., (1997) reported inferior parietal lobe activation as well as middle frontal gyrus activation, when the participants performed a visual task. Moreover, Basile et al., (1997) showed temporal and occipital sources of the target P3, with the centers of the neural activity in the vicinity of the superior temporal sulcus, in the hippocampal formation and parahippocampal gyrus and in the occipital extrastriate cortex (Basile et al., 1997, for a review see Herrmann & Knight, 2001). These are just a few examples of sources found so far when localizing the P3 component.

Considering the sheer number of P3m sources identified in various modalities and tasks, the exact identification of the sources in each specific task seems to be of great importance, when NIBS are considered to be applied as a tool for the modulation of single ERF components. Investigating the sources of task difficulty dependent P3m modulation additionally offers the opportunity to exactly target the sources of the modulation, which in turn could pave the way for clinical applications. Since, to the best of our knowledge, the sources of the task difficulty dependent P3m modulation was not localized previously, it is difficult to classify our results in the literature. However, considering the various source localization of the P3m, it is conceivable to assume that the source(s) of the task difficulty dependent P3m modulation varies between modalities and tasks.

## 7 Author Contributions

CB, FHK and CSH conceptualized the manuscript. CB programmed the experiment. CB and FHK collect data. CB wrote the draft manuscript. CB analyzed the data. All authors contributed to manuscript revision, read, and approved the submitted version.

